# Maternal obesity driven changes in collagen linearity of breast extracellular matrix induces invasive mammary epithelial cell phenotype

**DOI:** 10.1101/2022.11.29.518205

**Authors:** Jensen N. Amens, Gökhan Bahçecioglu, Kiera Dwyer, Xiaoshan S. Yue, M. Sharon Stack, Tyvette S. Hilliard, Pinar Zorlutuna

## Abstract

Obesity has been linked with numerous health issues as well as an increased risk of breast cancer. Although effects of direct obesity in patient outcomes is widely studied, effects of exposure to obesity-related systemic influences *in utero* has been overlooked. In this study, we investigated the effect of multigenerational obesity on epithelial cell migration and invasion using decellularized breast tissues explanted from normal female mouse pups from a diet induced multigenerational obesity mouse model. We first studied the effect of multigenerational diet on the mechanical properties, adipocyte size, and collagen structure of these mouse breast tissues, and then, examined the migration and invasion behavior of normal (KTB-21) and cancerous (MDA-MB-231) human mammary epithelial cells on the decellularized matrices from each diet group. Breast tissues of mice whose dams had been fed with high-fat diet exhibited larger adipocytes and thicker and curvier collagen fibers, but only slightly elevated elastic modulus and inflammatory cytokine levels. MDA-MB-231 cancer cell motility and invasion were significantly greater on the decellularized matrices from mice whose dams were fed with high-fat diet. A similar trend was observed with normal KTB-21 cells. Our results showed that the collagen curvature was the dominating factor on this enhanced motility and stretching the matrices to equalize the collagen fiber linearity of the matrices ameliorated the observed increase in cell migration and invasion in the mice that were exposed to a high-fat diet. Previous studies indicated an increase in serum leptin concentration for those children born to an obese mother. We generated extracellular matrices using primary fibroblasts exposed to various concentrations of leptin. This produced curvier ECM and increased breast cancer cell motility for cells seeded on the decellularized ECM generated with increasing leptin concentration. Our study shows that exposure to obesity *in utero* is influential in determining the extracellular matrix structure, and that the resultant changes in collagen curvature is a critical factor in regulating the migration and invasion of breast cancer cells.

## Introduction

The prevalence of obesity for U.S. adults in 2017 was 42.4%^1^. This is alarming, because obesity is listed by the Center for Disease Control (CDC) among the risk factors for cancer^2–4^ and is associated with poor prognosis and recurrence in breast cancer^5–7^. Obese postmenopausal women have a 20%-40% increased risk for developing breast cancer^4,8–10^.

Obesity has a systemic effect on the body. Immune cells in the adipose tissue of lean mice predominantly secrete anti-inflammatory cytokines, whereas in obese mice, the immune cells secrete proinflammatory cytokines^11^, indicating that obesity can lead to an increased inflammatory response^12^. which has been shown to be pro-tumorigenic^13^. Adipose tissue up-regulates collagens, fibronectin, matricellular proteins, and many extracellular matrix (ECM) remodeling enzymes^14^. These biochemical changes significantly influence the structure of the ECM, which, in turn, provide an environment that supports cancer initiation and progression^15–17^. For instance, obesity has been shown to promote the differentiation of adipocyte progenitor cells into myofibroblasts, which contributes to stiffening of the ECM^18^. Cancer cells are also known to block the differentiation of adipocyte progenitors to adipocytes on stiff substrates^19,20^, further supporting cancer progression.

Dietary factors and nutrition are known to affect breast cancer risk^21^, and a recent study showed that children of obese mothers also have increased risk of cancer^22^. The study conducted with 2 million children born between 2003 and 2016 showed that children born to severely obese mothers (body mass index > 40) had a 57% higher risk of developing leukemia before age 5 compared to children born to normal weight mothers, clearly showing that maternal obesity is also a risk factor for cancer. Many studies have been conducted investigating the effect of obesity on breast cancer and breast cancer survival. Given that in the USA almost 50% of new mothers are either overweight or obese^23^, and the obesity rates are increasing dramatically in recent years^1^, there is an urgent need for additional studies comparing *postpartum* vs. *in utero* exposure to obesity.

Here, we investigate the influence of multigenerational obesity on ECM properties such as stiffness, collagen structure, and biochemical changes, and the subsequent effect on normal and cancerous breast epithelial cell behavior. We used mammary glands of female mice fed either high or low-fat diets, and whose dams were also fed either high or low-fat diets during pregnancy. After decellularization of the mammary glands, we seeded normal (KTB-21) and cancerous (MDA-MB-231) human mammary epithelial cells and tested their migratory and invasive behavior to compare the effects of *in utero* exposure to obesity to *postpartum* exposure on the breast ECM. This study is the first to investigate the effect of multigenerational obesity on ECM properties, most importantly on collagen fiber curvature, and its influence on the regulation of normal and cancerous epithelial cell behavior.

## Materials and Methods

### Harvesting mouse mammary glands

C57BL/6J mice were handled in accordance with the IACUC with the approval of the University of Notre Dame, which has an approved Assurance of Compliance on file with the National Institutes of Health, Office of Laboratory Animal Welfare. Female C57BL/6J mice were fed either a high-fat diet (western diet containing 40% fat from corn oil and butter as the fat source; Research Diets D12079B) or low-fat diet (normal diet containing 10% fat from corn oil as the fat source; Research Diets 98121701). When confirmed obese (5-gram weight differential) one low-fat and one high-fat diet fed mouse (dam) was bred with the same male mouse (fed low-fat diet) to control for paternal influence. The female pups from each dam were subsequently randomized and fed either high or low-fat diets until the high-fat diet groups became overweight/obese (∼6-8 months) resulting in four female pup cohorts: high-fat dam, high-fat pup (HH); high-fat dam, low-fat pup (HL); low-fat dam, high-fat pup (LH); and low-fat dam, low-fat pup (LL) (Fig. 1A).

**Figure 1:**
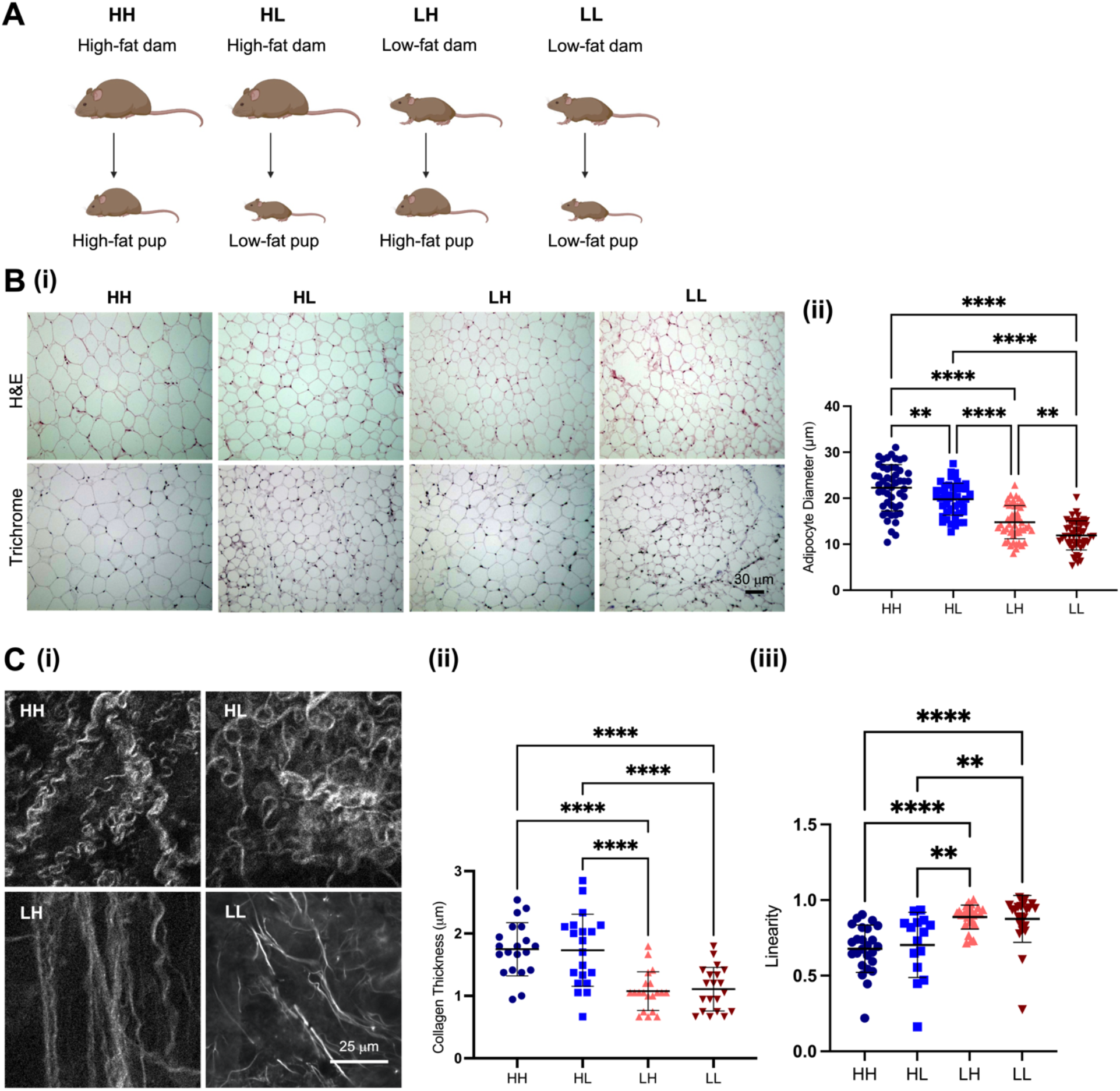
Characterization of model groups (A) Depiction of the four different model groups: HH, HL, LH, LL. (B) Adipocyte diameter of the model group tissues. i) Representative images showing hematoxylin and eosin (top) and Masson’s trichrome (bottom) staining of the tissues. n=2. ii) Quantification of the average adipocyte diameter from the histology images. (C) Collagen fiber structure in the matrices. i) Representative second harmonic generation images showing the collagen fibers. n=3. ii) Quantification of the average fiber diameter from the SHG images. iii) Linearity of the collagen fibers. For statistical significance, one-way ANOVA was performed for (B) and (C) followed by Tukey’s HSD, and two-way ANOVA for (B). **P<0.01, and ****P<0.0001.

Mice were euthanized and dissected, and their fourth mammary glands were collected. Tissues harvested were used immediately or wrapped in aluminum foil, flash frozen in liquid nitrogen, and stored at -80 °C until use.

### Decellularization

To section the tissues in the cryostat, tissues were thawed at room temperature, blotted on a tissue paper, embedded in Tissue-Tek optimum cutting temperature (O.C.T.) compound (Sakura, USA), frozen at -20 °C, and sectioned at 300 μm thickness. Sections were placed in PBS to remove the O.C.T compound.

Tissue decellularization was performed as described previously^16^. To remove cells, whole tissues or sections were incubated in 0.5% SDS for two days at 4 °C, with gentle agitation and SDS change every 12 hours. To remove the lipids, whole tissues or sections were incubated in 100% isopropanol for two days at 4 °C, with gentle agitation and isopropanol change every 12 hours. After decellularization, matrices were washed twice with PBS and stored at 4 °C until use.

### Mechanical characterization

For the mechanical characterization, a nanoindenter (Piuma Chiaro, Optics11, The Netherlands) with a 10 N load cell, and a silicon nitride SNL-10 cantilever (Bruker, USA) with a spring constant of around 0.261 N/m was used. The native tissues (n = 2) were tested at 5 different locations. The loading velocity was 2 mm/s. Young’s modulus was determined by a custom developed MATLAB code using Hertz contact model as described previously^16^, assuming a Poisson’s ratio of 0.5. For testing of decellularized, delipidized ECM, a probe with radius of 21 μm and spring constant of around 0.320 N/m was used. The smaller radius probe was used for the ECM as it had a higher spring constant for the stiffer ECM and the smaller radius allowed us to measure the stiffness of the specific collagen fibers within the ECM. The loading velocity was 2 mm/s. The ECM (n = 3) were tested at 10-20 different locations.

### Histology

Model group breast tissue samples were sectioned at 6 μm thickness and stained with hematoxylin and eosin (H&E) and trichrome at the Notre Dame Integrated Imaging Core Facility to analyze the adipocyte density and the adipocyte size of the samples. Adipocyte diameter was determined manually using ImageJ.

### Multiphoton microscopy

Decellularized sections were used for second harmonic generation (SHG) imaging using a Multiphoton microscope (Olympus, FV1000 MPE, USA) to analyze the collagen structure of the samples. ImageJ software (NIH, USA) was used to measure the thickness and linearity of the collagen fibers in the SHG images. For thickness, 10 fibers were analyzed per image and 5 images per sample (N=3). Linearity was measured as the ratio of the straight distance between the two ends of the collagen fiber to the actual length of this fiber. For linearity, 5 fibers were analyzed per image and 5 images per sample (N=3).

### Cytokine Profiling

To profile the cytokines in the mouse breast, whole native tissues (n=3 pooled samples for each model group) were flash frozen in liquid nitrogen and ground using a mortar and pestle, and the resulting powders were suspended in protease inhibitor cocktail (Sigma, Cat No: P8340) solution and homogenized using an ultrasonicator. Next, Triton X-100 was added (final concentration: 1%) to disrupt cells and fatty components. The mixture was centrifuged 5 times at 10,000g for 5 minutes, with removal of the cellular debris and fatty components after each centrifugation cycle. Protein quantification was done using the bicinchoninic acid (BCA) rapid gold assay kit (Pierce, Thermo Fisher Scientific).

The relative cytokine content (Figure S2) was determined using the dot blot-based mouse XL cytokine array kit for the model group breast tissues and human XL cytokine array kit and human XL oncology array kit for the human cell conditioned media following manufacturer’s instructions (R&D Systems, Cat. No: ARY028). Briefly, array membranes were blocked with an array buffer for 1h at RT, washed and incubated overnight at 4 °C in equal amounts of the tissue or cell media effluent extracts. After washing, the membranes were incubated in the biotinylated antibody cocktail solution for 1 h, in streptavidin-horseradish peroxidase (HRP) for 30 min, and in the chemiluminescent reagent for 1 min. Membranes were exposed to X-ray for 5-10 min using a biomolecular imager (ImageQuant LAS4000, GE Healthcare, USA). Relative cytokine content was determined after quantification of the dot intensity using ImageJ.

### Cell culture

MDA-MB-231 breast cancer cells tagged with a green fluorescent protein (GFP) reporter were a kind gift from Dr. Siyuan Zhang at University of Notre Dame. Cells were cultured in basal medium (DMEM high glucose medium supplemented with 10% fetal bovine serum (FBS) and 1% penicillin-streptomycin (P/S)). The culture was maintained with media changes every two days until 90% confluent. Cells were treated with trypsin, resuspended in basal media, counted, and seeded on the decellularized matrices.

KTB-21 normal mammary epithelial cells were kindly obtained from Dr. Harikrishna Nakshatri of Indiana University. Cells were cultured in basal medium (DMEM high glucose with F-12 medium supplemented with 10% fetal bovine serum (FBS) and 1% penicillin-streptomycin (P/S)). The culture was maintained with media changes every two days until 90% confluent. Cells were treated with trypsin, resuspended in basal media, counted, and seeded on the decellularized matrices.

Rat cardiac fibroblasts were isolated from 2-day-old neonatal rat pups and passaged to remove any differing cell types. Primary fibroblasts were then cultured in basal medium (DMEM high glucose medium supplemented with 10% fetal bovine serum (FBS) and 1% penicillin-streptomycin (P/S)).

Mouse mammary fibroblasts were purchased from Fisher Scientific. Cells were cultured in basal medium (Fibroblast media [Cell Biologics] supplemented with 10% fetal bovine serum (FBS), 1% antibiotic-antimycotic solution, 1% L-Glutamine, 0.1% hydrocortisone, and 0.1% fibroblast growth factor).

### Migration Assay

To study the cell motility and migration of MDA-MB-231 and KTB-21 cells within the matrix environment, cells were seeded onto the decellularized matrices at 10^5^ cells/matrix, and cultured for 24 hours in basal medium, and imaged under a fluorescence microscope (Zeiss Axio Observer.Z1) at 15 min intervals for six hours. The time-lapse images were then converted to .AVI video files and analyzed using the mTrackJ plug-in in the imageJ software and the cell motilities calculated.

Additionally, migration assays were done with stretched matrices. Briefly, the length of the matrices was measured, and matrices were stretched at 10% strain and pinned onto a disk-shaped polydimethylsiloxane (PDMS) mold to straighten the collagen fibers in the matrices. Cells were then seeded on the stretched matrices at a 2 × 10^5^ cells/matrix density, and imaged as described above.

### Invasion assay

MDA-MB-231 cells were seeded on the decellularized matrices and cultured for seven days in basal medium at 37°C. The seeded matrices were then placed into the transwell inserts against 15% FBS basal medium placed in the bottom well. After five days in the assay, the cells that invaded through the transwell inserts to the bottom wells were imaged using the fluorescence microscope and counted.

Alternatively, the invasion assays were done with the cell-seeded stretched matrices. The invaded cells were counted on day 5.

### Primary Fibroblast ECM Leptin Assay

18 mm diameter glass slides (VWR) were coated in a 25 mg/mL gelatin from porcine skin solution (Sigma Aldrich) and left to gel for one hour at room temperature. The slides were then placed in the bottom of 12-well plates and primary fibroblasts from rats or mice were seeded on the wells. The primary fibroblasts were cultured in basal medium until confluency. After confluency was reached, basal medium supplemented with varying leptin (Abcam) concentrations (5 ng/mL, 10 ng/mL, 20 ng/mL, 30 ng/mL, 40 ng/mL) replaced the original basal medium. Every other day this medium was replaced with basal medium supplemented with leptin and L-ascorbic acid (Sigma Aldrich) at a concentration of 50 μg/mL. This procedure was repeated for three weeks. After the three weeks, when enough ECM was produced by the primary fibroblasts, the ECM was decellularized using 0.5% SDS changed twice per day with gentle agitation. After decellularization, the ECMs were imaged using SHG and stained for collagen I and imaged before performing migration assays. SEM imaging was conducted using a gold palladium coating of 9 nm on the samples.

### Statistical Analysis

Data were analyzed for statistical significance using GraphPad Prism 6. One-way analysis of variance (ANOVA) followed by Tukey’s HSD or two-tailed Student’s t-test were performed to compare the results. Data are presented as the mean ± standard deviation (SD).

## Results

### Maternal diet influences breast ECM structure in offspring

The weights of the four female pup cohorts after 8-10 months on the respective diets were as in Table 1. As expected, pups that were fed a high-fat diet (HH and LH) weighed 22% more than those fed a low-fat diet (HL and LL; P<0.001), regardless of the dam diet. To analyze the adipocyte diameter and the lipid content of the tissue samples, we performed H&E and trichrome staining (Fig. 1B(i)). Quantification of the images revealed that adipocytes in the LL group (11.9 ± 3.2 μm) were significantly smaller than those in the other groups (P<0.01), confirming that the adipocytes in this group has less lipid content. Adipocytes in the HH group (22.3 ± 5.0 μm) were the largest (P<0.01), indicating a high lipid storage within these cells (Fig. 1B(ii)). The adipocytes in HL (19.8 ± 3.5 μm) and LH (14.8 ± 3.6 μm) had varying diameters, indicating variability in the lipid contents.

**Table 1.** Weights of the model group mice after 6-8 months of diet. n = 5 mice.

| Diet group | Weight (g) |
| --- | --- |
| HH | $28.0 \pm 2.1$ |
| HL | $21.8 \pm 1.9$ |
| LH | $29.0 \pm 3.6$ |
| LL | $22.6 \pm 1.3$ |

We then imaged the collagen in the matrices using SHG microscopy and analyzed the images using ImageJ to calculate collagen fiber thickness and linearity. Fiber thickness in the HH (1.02 ± 0.46 μm) and HL (1.21 ± 0.59 μm) matrices was higher than that in the LH (0.743 ± 0.31 μm) and LL (0.687 ± 0.32 μm) groups (P<0.0001) (Figs. 1C(i) and C(ii)). Additionally, collagen fibers in HH and HL groups were curvy and interconnected, while those in the other groups were straight (Fig. 1C(i)). Quantification of fiber linearity confirmed that collagen fibers in the LL and LH matrices were straighter than those in the HH and HL matrices (P<0.001) (Fig. 1C(iii)). These results highlight the predominant effect of maternal diet on offspring collagen properties. Stiffness testing was conducted on both breast tissue and decellularized, delipidized breast ECM using a nanoindenter (Fig. S1). The breast tissue did not display any significant differences between the stiffnesses of the model groups. The stiffness of the ECM displayed only significance between the LH group and the other model groups but displayed no consistent trend throughout the samples, therefore, we determined stiffness to not be a significant factor (Fig. S1B).

Adipocytes secrete several cytokines, known as adipokines, that regulate appetite and satiety, energy expenditure, inflammation, insulin sensitivity and energy metabolism^24^. Cytokine arrays were performed to characterize the adipokine and cytokine contents of the native mouse breast tissues. Overall, higher cytokine levels were found in the HH group compared to the other groups, although the results were not significant (Fig. S2A).

### Maternal diet has an impact on cell migration and cytokine production

After observing structural differences in the ECMs of the model group tissues, the next step was to investigate how these changes would affect breast cancer cell migration and invasion behavior. We seeded MDA-MB-231 cells on the decellularized matrices and tracked the migration of cells over a six-hour period. We found that cells were more motile on the matrices obtained from high-fat diet dams (Fig. 2C, Fig. S3). Quantification revealed that cell motility was significantly higher on the HH (2.25 ± 0.83 μm/h) and HL (2.13 ± 0.40 μm/h) groups than on the LH (1.23 ± 0.47 μm/h) and LL (0.87 ± 0.34 μm/h) groups (P<0.0001) (Fig. 2C). This same trend was also found when studying the migration of normal breast epithelial KTB-21 cells on the model group matrices (Fig. S4). The KTB-21 cells seeded on the HH, and HL matrices were significantly more motile than those seeded on the LH and LL matrices (P<0.0001). Since the secretion of cytokines can promote cancer progression and migration^25^, we investigated if differences in cytokine production for cells seeded on the model group matrices could be the cause for this increase in migration. While the cytokine production for cells seeded on the HH tissue was greater than that of the other model groups, the difference was not significant (Figs. S2B, S2C).

**Figure 2:**
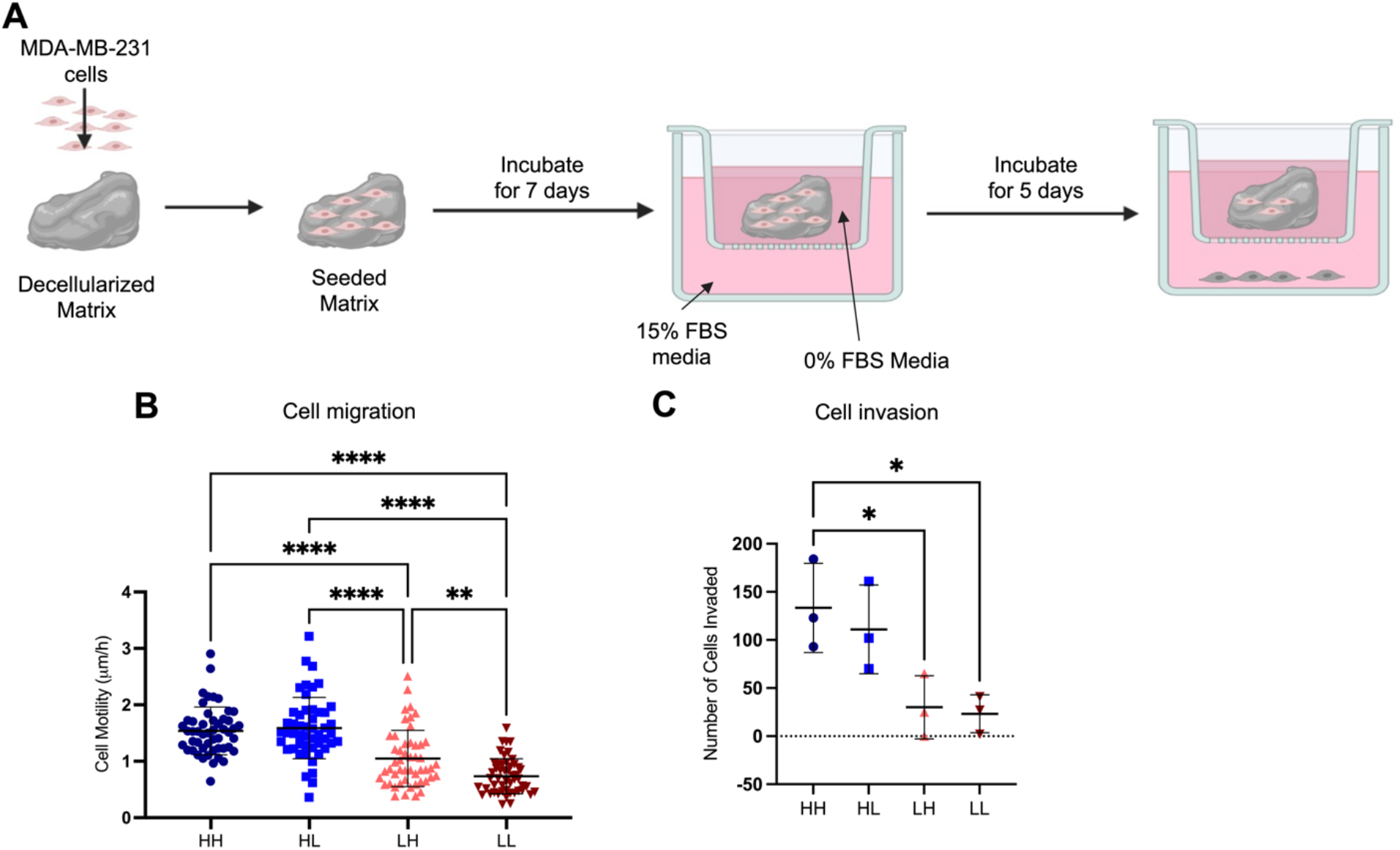
Migration and invasion MDA-MB-231 cells are increased on matrices obtained from mice with maternal high-fat diet. (A) Schematic showing the experimental procedure for the transwell invasion assay. (B) Average cell motility of MDA-MB-231 cells based on the fluorescence images over six hours n=2. (C) Number of cells invaded through each of the model group matrices n=3. For statistical significance, one-way ANOVA was performed for (B) and (C) followed by Tukey’s HSD. * denotes statistical significance with P<0.05, ** P<0.01, and **** P<0.0001.

### Maternal diet impacts breast cancer cell invasion

Next, we investigated the breast cancer cell invasion through matrices of the different diet groups (Fig. 2A). We saw an invasion pattern similar to that of the cell motility from the migration assay. Cell invasion was significantly higher through the HH and HL matrices (102.25 cells ± 63.04 cells and 91.25 cells ± 47.28 cells, respectively) than the LH and LL (54.5 cells ± 48.36 cells and 26.5 cells ± 15.01 cells, respectively) after five-day incubation in transwell inserts (P<0.0001) (Fig. 2B).

### Increasing collagen linearity decreases cell migration

As the collagen fiber linearity was significantly higher in the LH and LL groups than HH and HL, we investigated if stretching the matrices slightly to straighten the collagen fibers, would impact cell migration on the matrices. Stretching the matrices to a 10% strain resulted in a decrease in the collagen curvature (increase in linearity) of the HH and HL matrices (P<0.0001) (Fig. 3C). In control experiments, we determined whether the low strain rate (10 %) we used in this study affects the stiffnesses of the stretched matrices by testing the unstretched and stretched matrices using a nanoindenter to assess stiffness (Fig. S1). The results showed only a significant difference in the stiffnesses of the LH unstretched (15.1 kPa ± 8.40) and LH stretched (9.26 kPa ± 9.17) matrices, where the LH unstretched ECM was slightly stiffer than that of the stretched ECM. Based on these results, we concluded that, as expected, stretching the HH and HL ECM at a low strain rate does not significantly affect the stiffness of the matrix.

**Figure 3:**
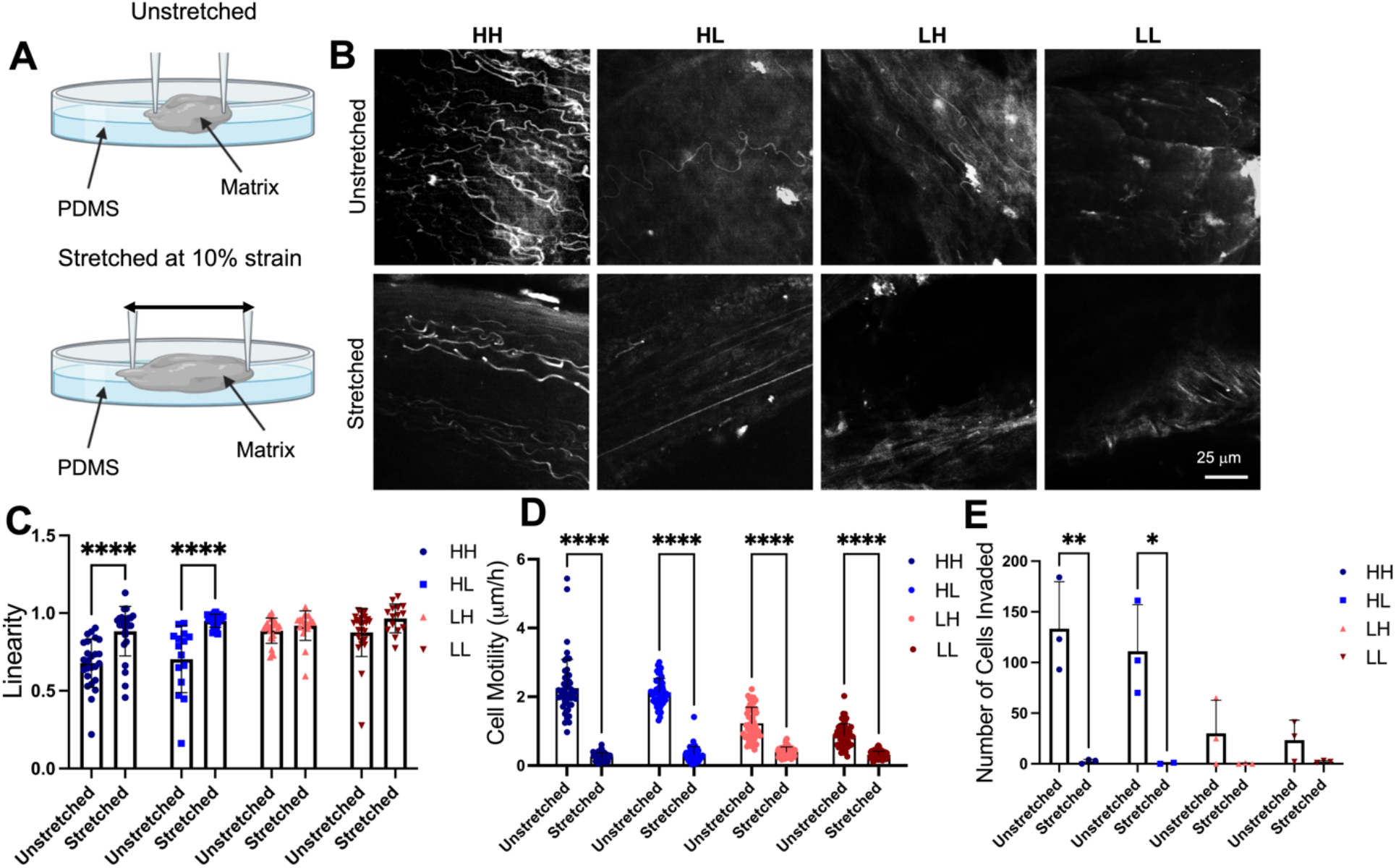
Stretching ECM significantly decreases the curviness of collagen fibers. (A) Schematic depicting the set-up for the stretching experiments. (B) SHG images of unstretched and stretched model group matrices. (C) Quantification of the linearity for the stretched and unstretched matrices. (D) Average cell motility of the stretched matrices based on the fluorescence images over six hours n=2. (E) Number of MDA-MB-231 cells invaded through each of the model group matrices after stretching n=3. For statistical significance, one-way ANOVA was performed for (C), (D), and (E) followed by Tukey’s HSD. **** denotes statistical significance with P<0.0001.

Motility of MDA-MB-231 cells on stretched sHH and sHL decreased significantly when seeded on stretched ECM (P<0.0001) (Fig. 3), showing that collagen fiber linearity is important in regulating cell migration speed. When compared to their unstretched counterparts, the cells seeded on the stretched (straightened, more linear) sHH and sHL matrices had significantly decreased motility (Fig. S5). The motility of cells seeded on the stretched sHH (0.257 μm/h ± 0.113) and sHL (0.332 μm/h ± 0.216) matrices were closer to, or even lower than, motility on the stretched sLH (0.396 μm/h ± 0.145) and sLL (0.310 μm/h ± 0.108) matrices (Fig. 3D). Compared to unstretched matrices, cell motility on the stretched, and therefore less curvy, matrices were significantly low. Additionally, this study was performed with the KTB-21 cells, which showed the same trend (Fig. S6). KTB-21 cells seeded on the stretched sHH and sHL matrices had significantly decreased motility when compared to their unstretched counterparts shown in figure S4.

### Collagen curvature increases invasion of breast cancer cells

We next examined the invasion of MDA-MB-231 cells on the stretched matrices. Stretching reduced the cell invasion considerably. Cell motility on the stretched HH and HL matrices was significantly lower than that on the unstretched matrices (P<0.038). After stretching, the number of cells invaded was the highest with the HL (6.33 cells ± 10.1 cells), and almost no invasion was detected in the other groups after five days of incubation in transwell inserts (Fig. 3E).

### Leptin effects the curvature of collagen produced by primary fibroblasts

Previous studies show an increase in leptin in the pups of obese dams^23,26,27^. We investigated the effect of leptin on the curvature of collagen produced by primary mammary fibroblasts in cell culture and subsequent motility for cancer cells seeded on the decellularized matrices produced by these fibroblasts. In mice, the serum leptin concentration in pups of obese dams (10-40 ng/mL) is significantly higher than that in pups of non-obese dams (5-20 ng/mL)^27^, therefore we used leptin concentrations of 5, 10, 20, 30, and 40 ng/mL to treat the fibroblasts. In our study, we found that an increase in the leptin concentration led to a significant decrease in collagen fiber linearity or an increase in curvature (P<0.0001), where the primary fibroblasts with no treatment led to a linearity of 0.98, and those treated with 20 ng/mL produced a linearity ratio of 0.96, those treated with 30 ng/mL leptin produced an ECM with a collagen fiber linearity of 0.74, and those treated with 40 ng/mL leptin produced collagen fibers with a linearity ratio of 0.81 (Fig. 4B). Remarkably, motility of MDA-MB-231 cells seeded on the decellularized form of primary fibroblast-generated matrices increased with the increase in leptin concentration and the resulting decrease in fiber linearity (Fig. 4C). The ECMs produced with 0 ng/mL and 20 ng/mL had significantly lower cell motilities of 0.81 μm/h ± 0.30 and 0.70 μm/h ± 0.29, whereas the ECM produced with 30 ng/mL leptin and 40 ng/mL had the highest average cell motilities of 2.23 μm/h ± 0.41 and 2.33 μm/h ± 1.1, respectively. We also investigated the effect of an increased leptin concentration on fibroblast differentiation to myofibroblasts. Increasing the leptin concentration within the media provided to the fibroblasts resulted in a significant increase in differentiation to myofibroblasts as shown through an increased staining for α-SMA (P<0.001) (Fig. S7).

**Figure 4:**
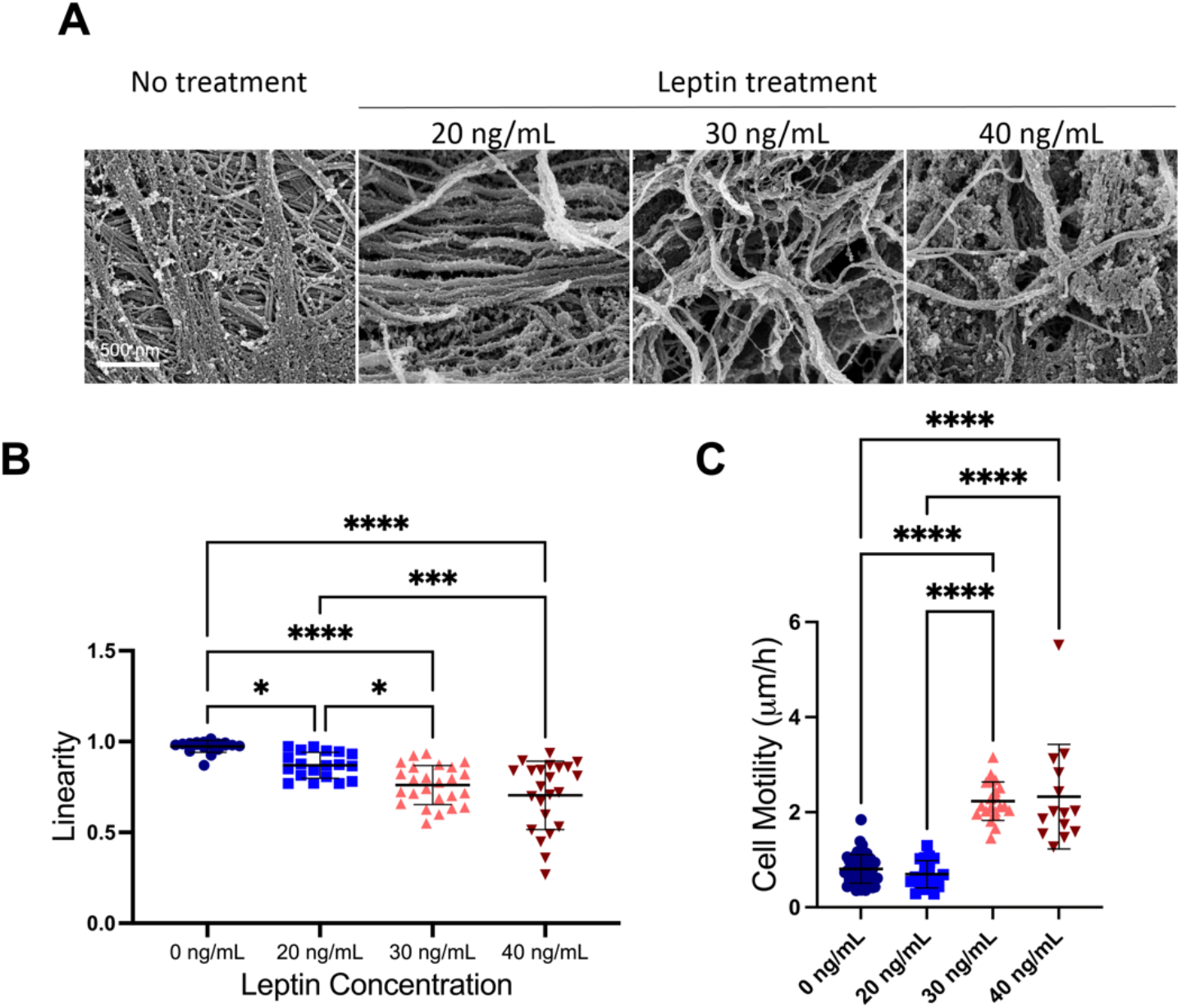
ECM generated by primary fibroblasts cultured with higher concentrations of leptin are curvier and increase cell motility. (A) SEM images of the ECM produced with the leptin concentration listed above the image. (B) Quantification of the linearity for the ECM generated with varying leptin concentrations, n = 2 (C) Average cell motility of MDA-MB-231 cells based on the fluorescence images over six hours, n = 3. For statistical significance, one-way ANOVA was performed for (B) and (C) followed by Tukey’s HSD. * denotes statistical significance with P<0.05, ***P<0.001, and **** P<0.0001.

### Cell motility correlates with fiber curvature and thickness but not with stiffness

Utilizing the results gathered above, we then compared fiber curvature, thickness, and stiffness each to the cell motility results to confirm the correlation between the characterizations. Cell motility had positive correlations with the fiber curvature (R^2^=0.89) and thickness (R^2^=0.93), but not with the matrix stiffness (R^2^=0.05) (Fig. S8).

## Discussion

Multiple studies have investigated the impact of obesity on cytokine production and how it influences the cancer microenvironment^14,18^. In this study, we specifically examined the alterations on breast ECM and collagen fiber structure due to multigenerational diet induced obesity and how these alterations in turn affect epithelial cell motility and invasion.

With the increased appreciation of the tumor microenvironment (TME) in tumor progression, the influence of obesity on tissue ECM and its effects on cancer cell phenotype have been of interest. In a recent study, Seo et al^18^ found an increase in linearity within the breast ECM collagen fibers for mice fed a high-fat diet compared to lean mice, which corresponds to the *postpartum* high-fat diet exposure group (LH) in our study. Our results demonstrated *postpartum* exposure to the high-fat diet alone (LH group) resulted in a slight, yet not significant, increase in linearity in the breast tissue collagen fibers as compared to the low-fat diet mice (LL group). Interestingly, we found a more profound effect of maternal diet on ECM properties relative to the diet of the pup itself. Breast tissues of pups exposed to a maternal high-fat diet (HH and HL) had significantly thicker and curvier collagen fibers, and larger adipocytes than those of mice exposed to a maternal low-fat diet, regardless of the pup’s diet *postpartum* (LH and LL). Cancerous (MDA-MB-231) and normal (KTB-21) human mammary epithelial cell migration and invasion were significantly increased when seeded on HH and HL matrices after decellularization, compared to cells seeded on LH and LL matrices. Because the mechanical properties and cytokine contents were not significantly different between the sample groups, we investigated the effect of the collagen fiber curvature on the migration and invasion of both KTB-21 and MDA-MB-231 cells by stretching the decellularized matrices prior to cell seeding to reduce the curvature and therefore increase the linearity. Straightening the collagen fibers decreased the migration and invasion of the cells seeded on the HH and HL matrices, without significantly changing the mechanical properties of the matrices. Finally, we tested the effect of leptin treatment on the structure of ECM deposited by primary fibroblasts to test our hypothesis that an increase in maternal blood leptin levels (corresponding to the leptin concentration within the medium) is the cause of the curvier collagen fibers we have observed in the HH and HL groups. Indeed, primary mammary fibroblasts produced significantly curvier collagen fibers in response to increased leptin concentration in their media in a dose-dependent manner. Furthermore, cancer cells seeded on the curvier ECM produced by cultured fibroblasts exposed to increasing leptin concentrations showed higher motility suggesting the *in utero* exposure to increased leptin concentrations due to maternal obesity might be the cause of ECM alterations we have observed in the pups born to high fat diet fed dams. Taken together our results suggests that maternal diet during pregnancy has a more marked effect on the ultrastructure of collagen in the mammary gland than does the diet of the pup itself, indicating a lasting effect of maternal obesity on this tissue.

We showed that the length of the path travelled by the cells and the cell speed on the curvier, more interconnected collagen fibers was greater than on the straight fibers. Furthermore, more cells invaded through the HH and HL matrices compared to the LL and LH ones. This is in line with studies that showed greater collagen mesh size and entangled fibers resulting in an increase in the motility of the cells because of the increased fiber density^28,29^. The curvier ECM of the HH and HL mice closely resembles the fibrotic tissue microenvironment, which would allow the breast cancer cells to have increased migration and invasion on the interconnected, fibrotic collagen^30,31^. Some other studies have reported an increase in directional cell migration on more aligned collagen fibers^32^. However, the higher cell migration on the aligned and straight fibers have been shown to be due to the efficient directional migration, and not due to an increase in the speed of the cells^33^. Here, we showed the speed of cells was higher on the curvier and denser collagen mesh.

The thickness of collagen fibers has also been reported to increase the cell motility of many different cell types, including breast cancer cells^28,34^, and our study showed an increase in collagen fiber thickness for the mice with an *in utero* exposure to obesity (HH and HL). This suggests another mechanism through which *in utero* exposure to obesity effects the structure of the ECM, in turn, increasing the migration and invasion of the breast cancer cells. Although the effect of obesity on the collagen thickness in the breast tissue has not been studied, the fiber diameter of the tendon collagen was shown to be larger in obese mice as compared to the lean mice^35^, confirming our analysis of the increased collagen fiber thickness within the ECM of the HH and HL model groups. Additionally, stiffness is known to influence cell migration^36,37^. However, in our study we found that high-fat diet has no significant effect on the stiffness of the breast tissues or matrices. Cell motility has good correlations with the fiber curvature and thickness, but not with the matrix stiffness. Thus, our results strongly indicate that collagen fiber curvature and thickness might be one of the major parameters playing a role in the enhanced cell migration and invasion on the HH and HL matrices compared to LH and LL matrices. Since it has been more closely associated with tumor associated collagen signature (TACS), a well-established theory within breast cancer literature that describes the organization of collagen in the TME, we focused on investigating the role of collagen curvature.

In order to test our hypothesis that curvier collagen fibers lead to increased cell migration and invasion, we stretched the matrices slightly to make the fibers more linear and monitored the motility and invasion of cancerous and normal epithelial cells on the matrices. We observed that cells seeded on these stretched matrices showed decreased migration and invasion compared to the unstretched matrices, despite the decreased curvature and increased alignment upon stretching. TACS describes how increased collagen structure linearity and alignment within the TME leads to tumor progression^38,39^. Although our results seem to conflict with established TACS literature, TACS characterizes the collagen structure of the tumor itself, whereas we characterized the normal breast tissue, and as such there is a cadre of collagen-associated proteins that differ between the breast ECM and the tumor ECM^40,41^. Our observation in this study is related to microenvironmental conditions and its influence on the earlier stages of cancer, and it indicates there might be stage dependent differences in different phases of cancer. The permissive collagen structure we observed in the obese dam groups in this study is also in line with our earlier observations on healthy aged breast ECM, another high-risk group for breast cancer development, which also showed curlier and denser collagen that enhanced both normal and cancerous epithelial cell migration and invasion^16^. As such, while linear collagen is more predominant in later stages of cancer and might play a key role in malignancy, curlier and denser collagen might be more influential in cancer initiation.

In addition, since we used only a 10% strain rate to stretch the matrices and confirmed experimentally that this low degree of strain does not affect the overall stiffness of the ECM^42^, we can confidently say that we were within the toe region of the stress strain curve of biologic materials^42^ as the matrices were not stretched to resistance and the strain did not significantly alter the stiffness of the matrices. As such, we do not believe that we have significantly altered the matrix stiffness or structure due to realignment of the fibers and breakage of crosslinks. Thus, our results strongly indicate that collagen fiber curvature might be one of the major parameters playing a role in the higher cell migration and invasion on the HH and HL matrices compared to LH and LL matrices.

High *in utero* leptin exposure could be the reason for the increased collagen curvature that we have observed in HH and HL matrices. Previous studies have shown that children born to obese mothers have a higher risk of developing leukemia and other childhood cancers, and larger birth size has been associated with increased risk of cancer^22^. It has been shown that murine offspring exposed to a maternal high-fat diet before birth have high levels of blood leptin^27^. White et al. have also shown that maternal obesity is necessary for observation of high-fat diet effects in their offspring^26^. In addition, increased leptin levels have been shown to increase collagen production^43,44^. The higher collagen production may lead to denser collagen mesh and, thus, curvier collagen fibers within the breast ECM of the pups with a high-fat dam (HH and HL). Our results demonstrate that exposure of cultured mammary fibroblasts to increased leptin levels resulted in deposition of ECM with significantly curvier collagen fibers, which is supported through previous studies^43,44^. Another study has shown that leptin levels are increased in pups of obese dams with similar concentrations to those tested within our study, further confirming our results^27^. It has been shown in mice with loss of function mutation in the gene encoding leptin hormone that the linearity of collagen fibers in the breast tissue increases^18^. We also verified that cells seeded on the decellularized matrix produced by these fibroblasts showed a higher motility similar to what was observed with our model group matrices. Leptin can also lead to a higher myofibroblast differentiation in pups from the maternal high-fat diet cohort. We confirmed this by treating primary mouse mammary fibroblasts with various leptin concentrations and staining for α-SMA to confirm myofibroblast differentiation. With increased leptin concentration, we observed an increase in the α-SMA staining, indicating increased myofibroblast differentiation. Myofibroblasts are known to deposit a more fibrotic ECM ^18,43–48^. Overall, the increase in leptin due to the exposure to *in utero* obesity may cause a decrease in collagen fiber linearity within the HH and HL model group ECM. Lipid content and adipocyte diameter may also impact the collagen fiber structure. High lipid content and larger adipocytes may lead to strain on the surrounding ECM. An increased mechanical strain on the tissues has been proven to result in curvier collagen^36^. A greater mechanical strain on the breast of the obese mice due to high fat content can explain the collagen curvature within the HH and HL samples.

Here, we show that the dam high fat diet during pregnancy is more important in regulating the collagen structure of her female offspring’s breast tissue than the offspring’s own high fat diet after birth. Mouse pups exposed to high-fat diet *in utero* develop significantly curvier and thicker collagen fibers than those exposed to low-fat diet *in utero*, while high fat diet after birth does not significantly affect collagen fiber structure. We also showed that *in utero* exposure to high levels of leptin is at least partially responsible for the curvy collagen production, and the curvy collagen leads to increased motility and invasion of normal and cancerous human mammary epithelial cells. These findings indicate that avoiding obesity during pregnancy could significantly reduce the offspring’s risk of developing metastatic breast cancer in the future, as parental medical history is an important risk factor for the patients’ health. Another implication of our findings is that leptin could be a target during pregnancy to prevent future metastatic breast cancer risk. Looking at breast cancer through the multigenerational obesity lens could allow for more accurate prognosis and better treatments for breast cancer.

## Supporting information

Title Page

Supplemental Images

## Acknowledgments

This study was funded by NIH award number 5R01EB027660-02 (P.Z.), K01 CA218305-01 (T.S.H.), RO1CA109545 (M.S.S.) and Walther Cancer Foundation, Harper Cancer Research Institute Cancer Cure Ventures Award number 0184.01. We would also like to acknowledge Dr. Siyuan Zhang for the gift of the MDA-MB-231 cells used in this study and Dr. Harikrishna Nakshatri of Indiana University for the gift of the KTB-21 cells used in this study. This work partially supported by the Notre Dame Integrated Imaging Facility.

## Data Availability Statement

The raw data required to reproduce these findings are available upon request. The processed data required to reproduce these findings are available upon request.

