## Supplemental Images for "Maternal obesity driven changes in collagen linearity of breast extracellular matrix induces invasive mammary epithelial cell phenotype"

**Other supplementary materials for this manuscript include the following:**

Movies S1 to S20

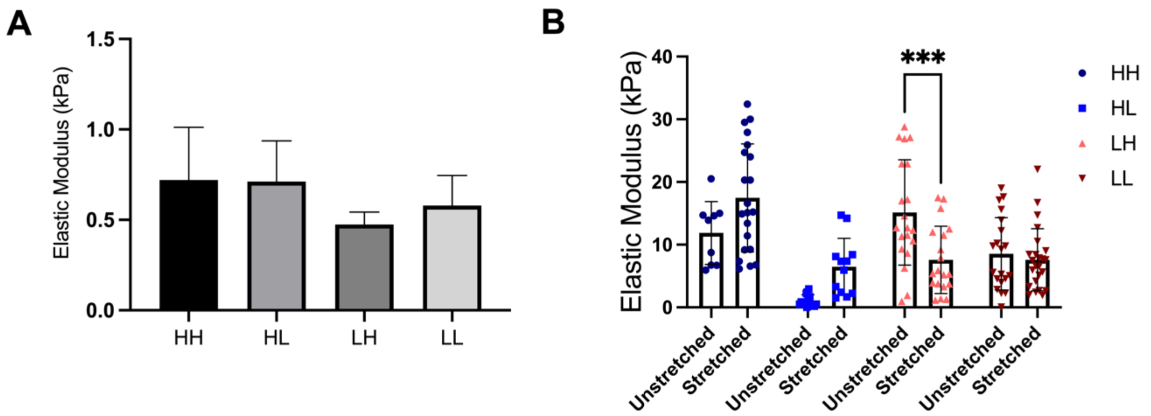


**Supplementary Figure 1:** Stiffnesses of model group tissues and ECM as measured by nanoindenting. (A) Stiffness measured using whole mouse breast tissue from each model group. (B) Stiffness measured using decellularized, delipidized ECMs from each model group in their unstretched and stretched (at 10% strain) forms. *** denotes P<0.001.

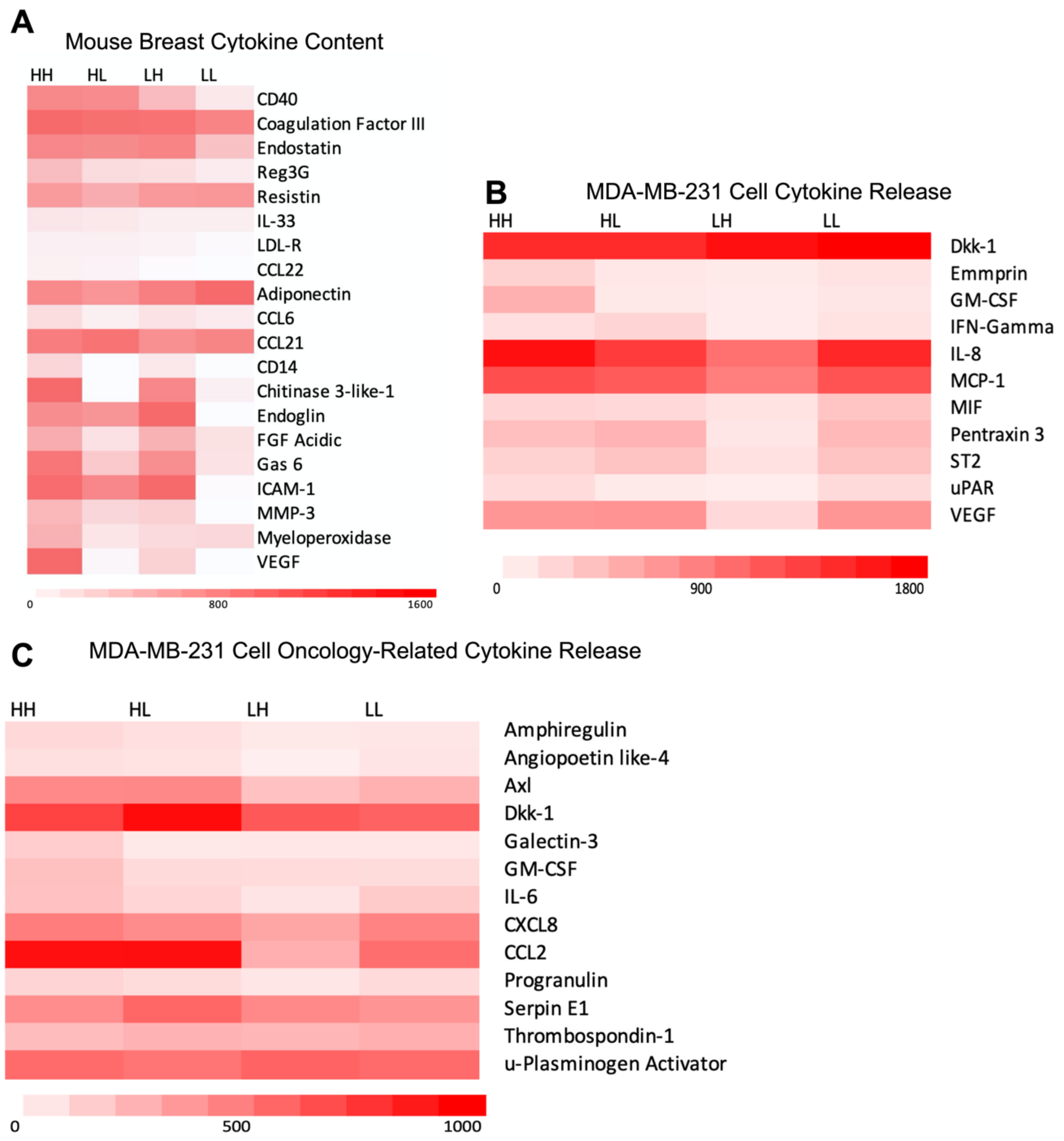


**Supplementary Figure 2:** Cytokine present in the mouse tissues and those produced by MDA-MB-231 cells on the decellularized matrices. Heat maps showing the (A) cytokines in the four model group mouse tissues. Heat maps showing the (B) cytokines and (C) cancer-related proteins released by MDA-MB-231 cells to cell culture media. Three samples were used for each group with two biological replicates.

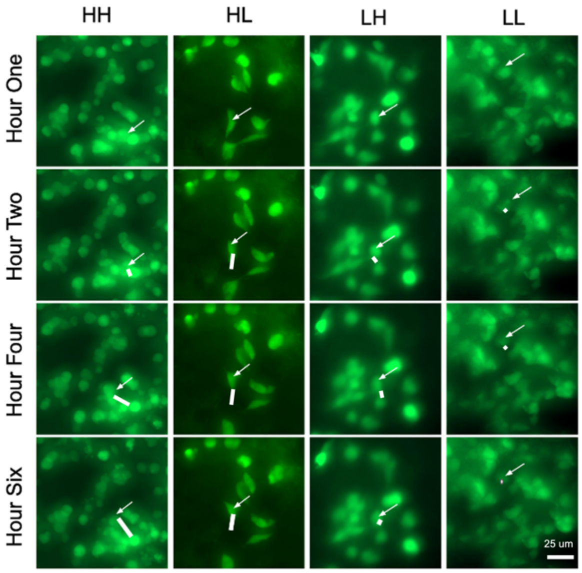


**Supplementary Figure 3:** (A) Fluorescence images of MDA-MB-231 cells seeded on each of the model group matrices. The white arrows track a cell through the images over time. The white bar in the Hour Six image is the distance the cell traveled over the course of the imaging.

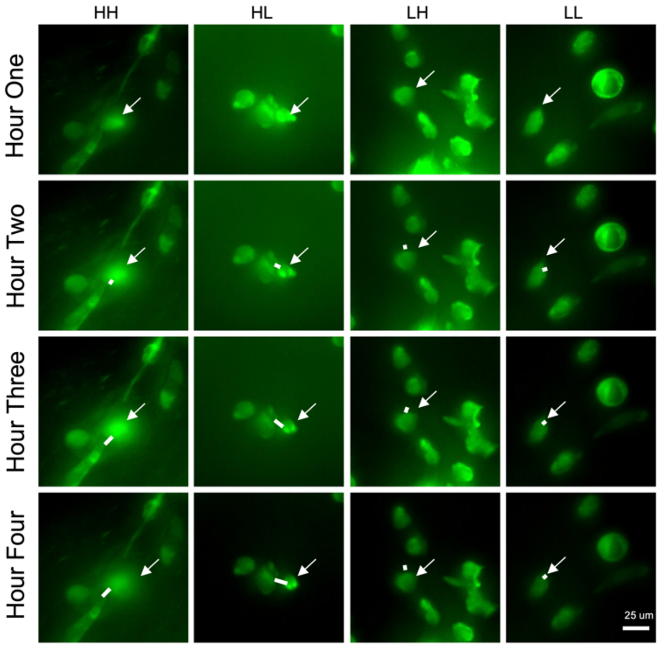


**Supplementary Figure 4:** (A) Fluorescence images of KTB-21 cells seeded on each of the model group matrices. The white arrows track a cell through the images over time. The white bar in the Hour Six image is the distance the cell traveled over the course of the imaging.

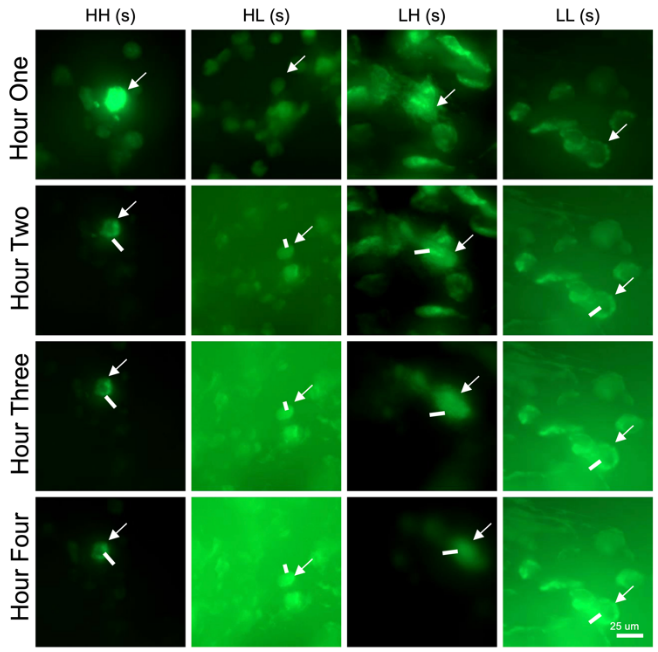


**Supplementary Figure 5:** Fluorescence images of MDA-MB-231 cells seeded on each of the stretched model group matrices. The white arrows track a cell through the images over time. The white bar in the image is the distance the cell traveled over the course of the imaging.

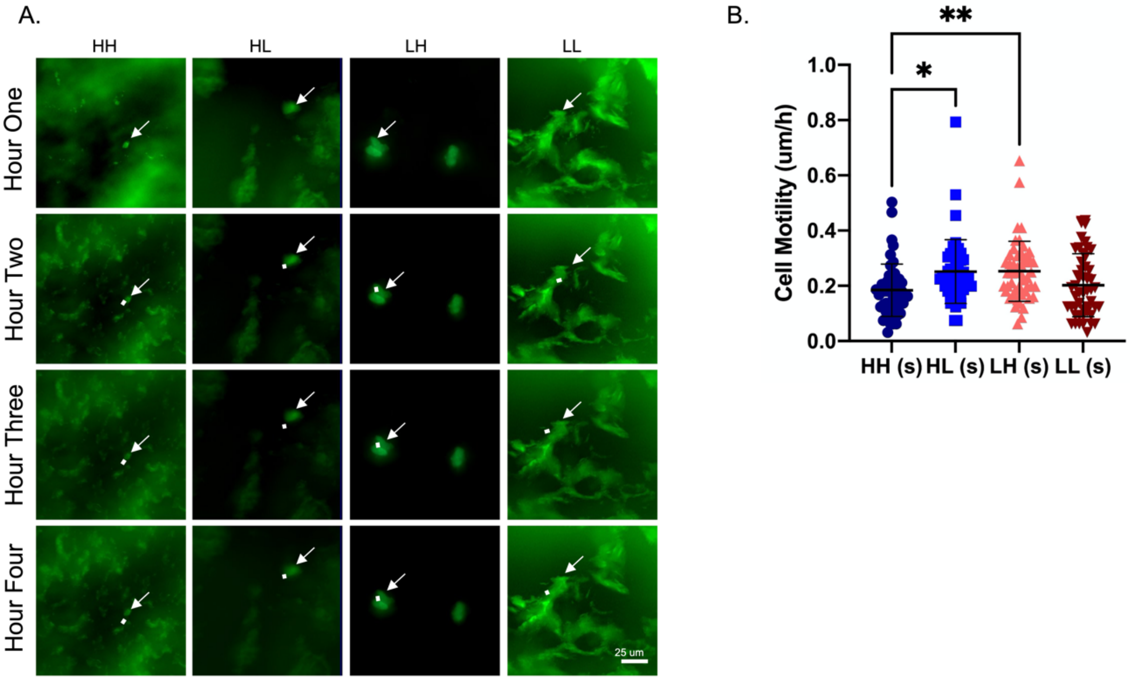

**Supplementary Figure 6:** Migration is decreased for KTB-21 cells seeded on stretched matrices. (A) Fluorescence images of cells seeded on each of the stretched model group matrices. The white arrows track a cell through the images over time. The white bar in the image is the distance the cell traveled over the course of the imaging. (B) Average cell motility based on the fluorescence images over six hours. For statistical significance, one-way ANOVA was performed for (B) followed by Tukey's HSD. * denotes statistical significance with P<0.05 and ** P<0.01.

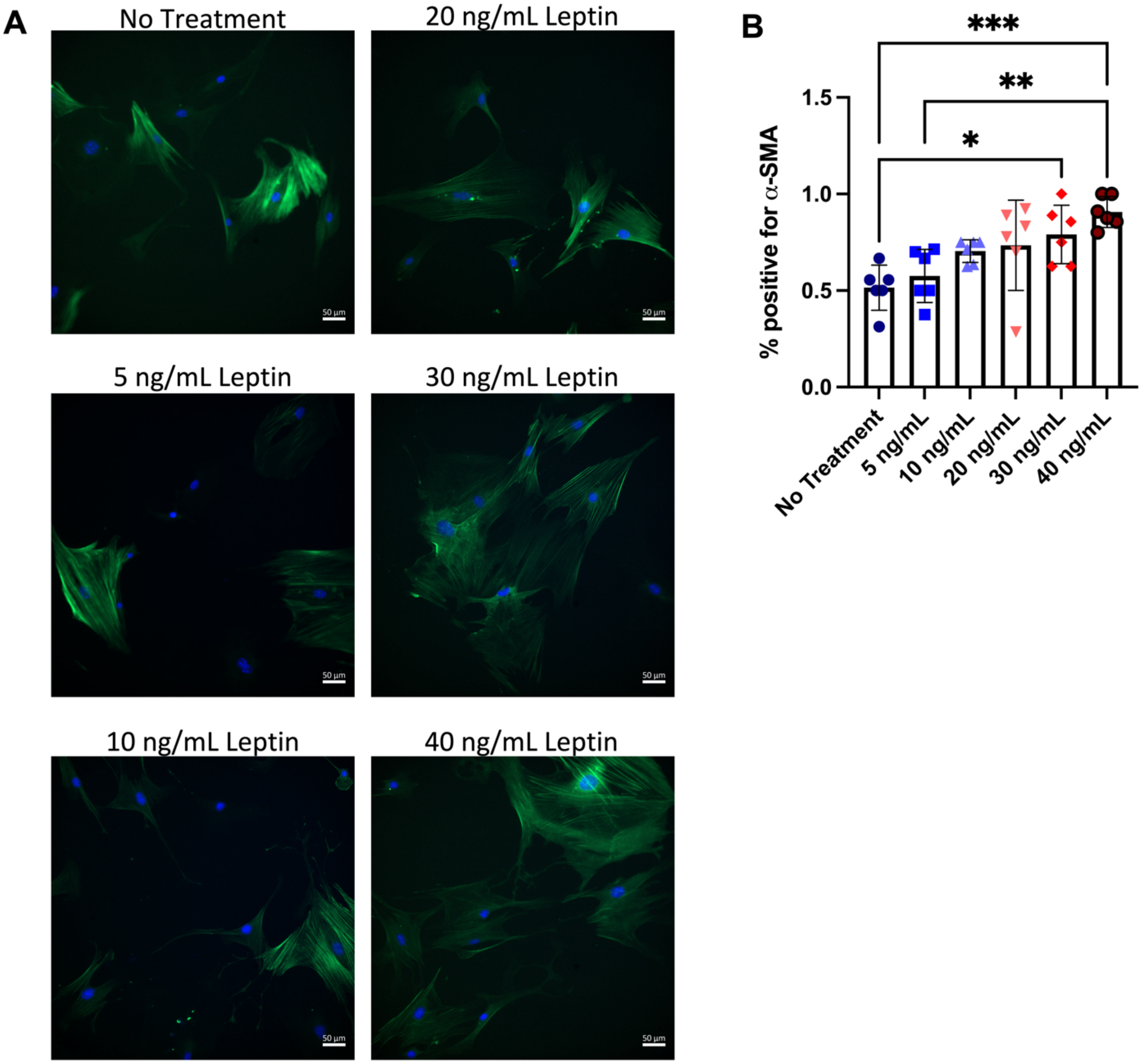

**Supplementary Figure 7:** α-SMA immunostain is increased in primary mouse mammary fibroblasts with increasing leptin concentration. (A) Fluorescence images of cells seeded on tissue culture plastic with α-SMA stained with green and DAPI stained with blue. (B) Quantification of cells positive for α-SMA based on fluorescence images. * denotes statistical significance with $P<0.05$, ** $P<0.01$, and ***$P<0.001$.

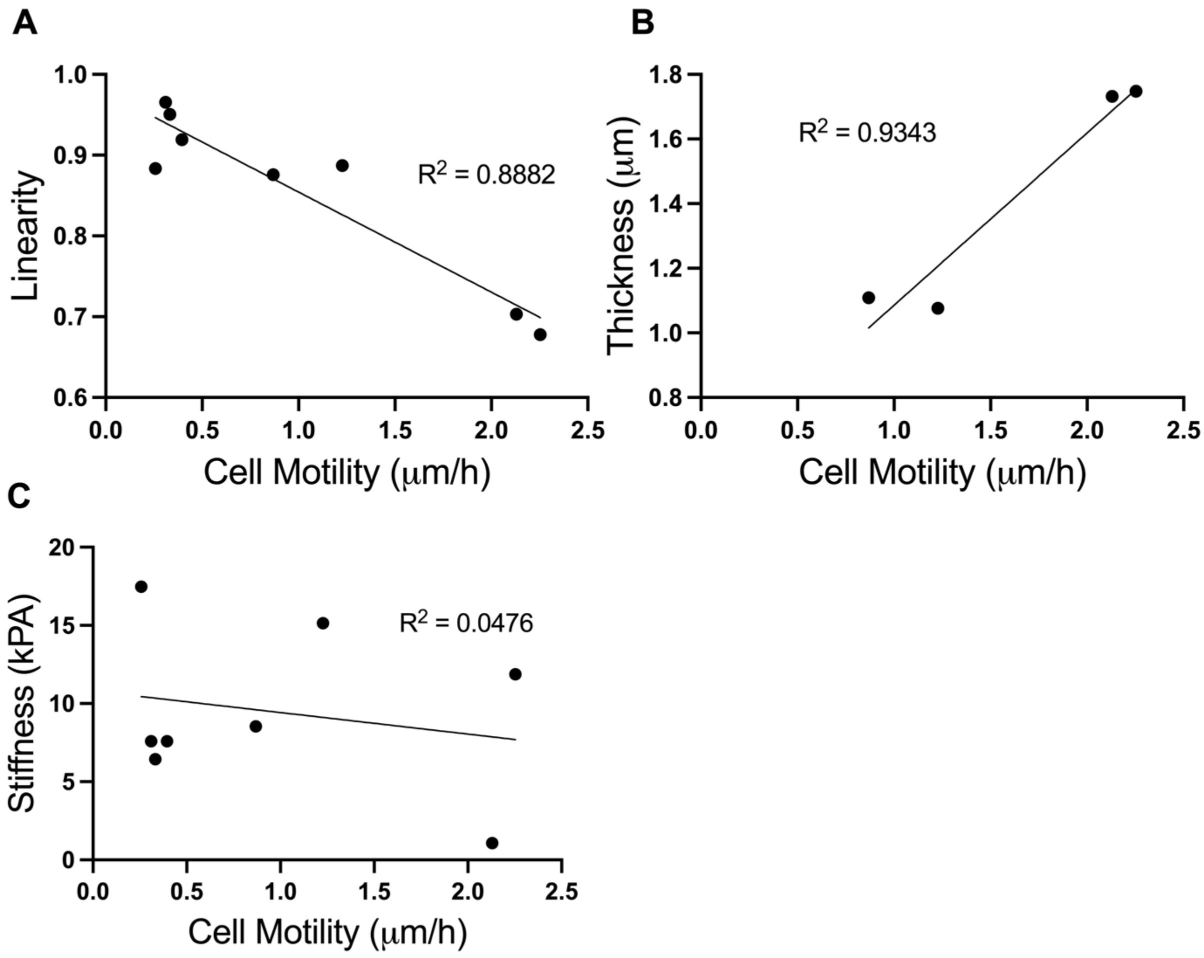


**Supplementary Figure 8:** Collagen curvature and thickness are positively correlated with cell motility, but stiffness is not correlated with cell motility. (A) Graph of Cell motility vs. Linearity with a linear trendline. (B) Graph of Cell motility vs. Thickness with a linear trendline. (C) Graph of Cell motility vs. Stiffness with a linear trendline.

**Table S1.** Mouse cytokines tested in dot blot assay of model group tissues

| Spot Location | Protein | Spot Location | Protein | Spot Location | Protein |
|---|---|---|---|---|---|
| **A1, A2** | Reference Spots | **D15, D16** | Cd26 | **G21, G22** | Il-33 |
| **A3, A4** | Adiponectin | **D17, D18** | Egf Endoglin | **G23, G24** | Ldlr |
| **A5, A6** | Amphiregulin | **D19, D20** | Endoglin | **H1, H2** | Leptin |
| **A7, A8** | Angiopoietin-1 | **D21, D22** | Endostatin | **H3, H4** | Lif |
| **A9, A10** | Angiopoietin-2 | **D23, D24** | Fetuin A (Ahsg) | **H5, H6** | Lipocalin-2 |
| **A11, A12** | Angiopoietin-like 3 | **E1, E2** | Fgf-a | **H7, H8** | Lix |
| **A13, A14** | Baff | **E3, E4** | Fgf21 | **H9, H10** | M-Csf |
| **A15, A16** | Cd93 | **E5, E6** | Flt-3 ligand | **H11, H12** | Mmp2 |
| **A17, A18** | Ccl2 | **E7, E8** | Gas 6 | **H13, H14** | Mmp3 |
| **A19, A20** | Ccl3 | **E9, E10** | G-CSF | **H15, H16** | Mmp9 |
| **A21, A22** | Ccl5 | **E11, E12** | Gdf15 | **H17, H18** | Myeloperoxidase (Mpo) |
| **A23, A24** | Reference Spots | **E13, E14** | Gm-Csf | **H19, H20** | Osteopontin (Opn) |
| **B3, B4** | Ccl6 | **E15, E16** | Hgf | **H21, H22** | Osteoprotegerin |
| **B5, B6** | Ccl11 | **E17, E18** | Icam-1 | **H23, H24** | Pd-Ecgf |
| **B7, B8** | Ccl12 | **E19, E20** | Ifn- $\gamma$ | **I1, I2** | Pdgf-bb |
| **B9, B10** | Ccl17 | **E21, E22** | Igfbp1 | **I3, I4** | Pentraxin 2 |
| **B11, B12** | Ccl19 | **E23, E24** | Igfbp2 | **I5, I6** | Pentraxin 3 |
| **B13, B14** | Ccl20 | **F1, F2** | Igfbp3 | **I7, I8** | Periostin |
| **B15, B16** | Ccl21 | **F3, F4** | Igfbp5 | **I9, I10** | Pref1 |
| **B17, B18** | Ccl22 | **F5, F6** | Igfbp6 | **I11, I12** | Proligerin |
| **B19, B20** | Cd14 | **F7, F8** | Il-1$\alpha$ | **I13, I14** | Pcsk9 |
| **B21, B22** | Cd40 | **F9, F10** | Il-1$\beta$ | **I15, I16** | Rage |
| **C3, C4** | Cd160 | **F11, F12** | Il-1ra | **I17, I18** | Rbp4 |
| **C5, C6** | Chemerin | **F13, F14** | Il-2 | **I19, I20** | Reg3G |
| **C7, C8** | Chitinase 3-like 1 | **F15, F16** | Il-3 | **I21, I22** | Resistin |
| **C9, C10** | Tissue factor | **F17, F18** | Il-4 | **I23, I24** | Reference Spots |
| **C11, C12** | Complement Component C5/C5a | **F19, F20** | Il-5 | **J1, J2** | Reference Spots |
| **C13, C14** | Complement factor D | **F21, F22** | Il-6 | **J3, J4** | E-Selectin |

| | | | | | |
|---|---|---|---|---|---|
| **C15, C16** | C-reactive protein | **F23, F24** | Il-7 | **J5, J6** | P-Selectin |
| **C17, C18** | Cx3cl1 | **G1, G2** | Il-10 | **J7, J8** | SerpinE1 (PAI- 1) |
| **C19, C20** | Cxcl1 | **G3, G4** | Il-11 | **J9, J10** | SerpinF1 |
| **C21, C22** | Cxcl2 | **G5, G6** | Il-12 p40 | **J11, J12** | Thrombopoietin |
| **D1, D2** | Cxcl9 | **G7, G8** | Il-13 | **J13, J14** | Tim-1 |
| **D3, D4** | Cxcl10 | **G9, G10** | Il-15 | **J15, J16** | Tnfα |
| **D5, D6** | Cxcl11 | **G11, G12** | Il-17A | **J17, J18** | Vcam-1 |
| **D7, D8** | Cxcl13 | **G13, G14** | Il-22 | **J19, J20** | Vegf |
| **D9, D10** | Cxcl16 | **G15, G16** | Il-23 | **J21, J22** | Wisp-1 |
| **D11, D12** | Cystatin C | **G17, G18** | Il-27 p28 | **J23, J24** | Negative Control |
| **D13, D14** | Dkk-1 | **G19, G20** | Il-28A/B | | |

**Table S2.** Human cytokines tested in dot blot assay of MDA-MB 231 cells seeded on matrices

| **Spot Location** | **Protein** | **Spot Location** | **Protein** | **Spot Location** | **Protein** |
|---|---|---|---|---|---|
| **A1, A2** | Reference Spots | **D11, D12** | IGFBP2 | **G13, G14** | MIF |
| **A3, A4** | Adiponectin | **D13, D14** | IGFBP3 | **G15, G16** | MIG |
| **A5, A6** | Apolipoprotein A-I | **D15, D16** | IL1α | **G17, G18** | MIP1α/MIP1β |
| **A7, A8** | Angiogenin | **D17, D18** | IL1β | **G19, G20** | MIP3α |
| **A9, A10** | Angiopoietin-1 | **D19, D20** | IL1ra | **G21, G22** | MIP3β |
| **A11, A12** | Angiopoietin-2 | **D21, D22** | IL2 | **G23, G24** | MMP9 |
| **A13, A14** | BAFF | **D23, D24** | IL3 | **H1, H2** | Myeloperoxidase |
| **A15, A16** | BDNF | **E1, E2** | IL4 | **H3, H4** | Osteopontin (OPN) |
| **A17, A18** | Complement component C5/C5a | **E3, E4** | IL5 | **H5, H6** | PDGF-AA |
| **A19, A20** | CD14 | **E5, E6** | IL6 | **H7, H8** | PDGF-AB/BB |
| **A21, A22** | CD30 | **E7, E8** | IL8 | **H9, H10** | Pentraxin 3 |
| **A23, A24** | Reference Spots | **E9, E10** | IL10 | **H11, H12** | PF4 |
| **B3, B4** | CD40 ligand | **E11, E12** | IL11 | **H13, H14** | RAGE |
| **B5, B6** | Chitinase 3-like 1 | **E13, E14** | IL12 p70 | **H15, H16** | RANTES |
| **B7, B8** | Complement factor D | **E15, E16** | IL13 | **H17, H18** | RBP4 |
| **B9, B10** | C-reactive protein | **E17, E18** | IL15 | **H19, H20** | Relaxin 2 |
| **B11, B12** | Crypto-1 | **E19, E20** | IL16 | **H21, H22** | Resistin |
| **B13, B14** | Cystatin C | **E21, E22** | IL17A | **H23, H24** | SDF1α |
| **B15, B16** | DKK-1 | **E23, E24** | IL18Bpa | **I1, I2** | SERPINE1 |
| **B17, B18** | DPPIV | **F1, F2** | IL19 | **I3, I4** | SHBG |
| **B19, B20** | EGF | **F3, F4** | IL22 | **I5, I6** | ST2 |
| **B21, B22** | EMMPRIN | **F5, F6** | IL23 | **I7, I8** | TARC |

| | | | | | |
|---|---|---|---|---|---|
| **C3, C4** | ENA-78 | **F7, F8** | IL24 | **I9, I10** | TFF3 |
| **C5, C6** | Endolgin | **F9, F10** | IL27 | **I11, I12** | TfR |
| **C7, C8** | FAS ligand | **F11, F12** | IL31 | **I13, I14** | TGFα |
| **C9, C10** | FGFb | **F13, F14** | IL32 | **I15, I16** | Thrombospondin 1 |
| **C11, C12** | FGF7 | **F15, F16** | IL33 | **I17, I18** | TNFα |
| **C13, C14** | FGF19 | **F17, F18** | IL34 | **I19, I20** | uPAR |
| **C15, C16** | FLT3 ligand | **F19, F20** | IP10 | **I21, I22** | VEGF |
| **C17, C18** | G-CSF | **F21, F22** | I-TAC | **I23, I24** | Reference Spots |
| **C19, C20** | GDF15 | **F23, F24** | Kallikrein 3 | **J1, J2** | Reference Spots |
| **C21, C22** | GM-CSF | **G1, G2** | Leptin | **J3, J4** | Vitamin D BP |
| **D1, D2** | GROα | **G3, G4** | LIF | **J5, J6** | CD31 |
| **D3, D4** | Growth Hormone | **G5, G6** | Lipocalin 2 | **J7, J8** | TIM3 |
| **D5, D6** | HGF | **G7, G8** | MCP1 | **J9, J10** | VCAM1 |
| **D7, D8** | ICAM-1 | **G9, G10** | MCP3 | **J23, J24** | Negative Control |
| **D9, D10** | IFN-γ | **G11, G12** | M-CSF | | |

**Table S5.** Human cancer-related proteins tested in dot blot assay of MDA-MB 231 cells seeded on matrices

| **Spot Location** | **Protein** | **Spot Location** | **Protein** | **Spot Location** | **Protein** |
|---|---|---|---|---|---|
| **A1, A2** | Reference Spots | **C15, C16** | HER2 | **F5, F6** | MMP2 |
| **A3, A4** | α-Fetoprotein | **C17, C18** | HER3 | **F7, F8** | MMP3 |
| **A5, A6** | Amphiregulin | **C19, C20** | HER4 | **F9, F10** | MMP9 |
| **A7, A8** | Angiopoietin-1 | **C21, C22** | FGFb | **F11, F12** | MST1 |
| **A9, A10** | Angiopoietin-like 4 | **C23, C24** | - | **F13, F14** | MUC1 |
| **A11, A12** | ENPP2 | **D1, D2** | MFH1 | **F15, F16** | Nectin 4 |
| **A13, A14** | AXL | **D3, D4** | FKHR | **F17, F18** | Osteopontin |
| **A15, A16** | BCL2L1 | **D5, D6** | Galectin 3 (GAL3) | **F19, F20** | TP27/KIP1 |
| **A17, A18** | CA125 | **D7, D8** | GM-CSF | **F21, F22** | TP53 |
| **A19, A20** | E-Cadherin | **D9, D10** | HCG | **F23, F24** | PDGF-AA |
| **A21, A22** | VE-Cadherin | **D11, D12** | HGF R | **G1, G2** | CD31 |
| **A23, A24** | Reference Spots | **D13, D14** | HIF1α | **G3, G4** | Progesteron R |
| **B1, B2** | - | **D15, D16** | HNF3β | **G5, G6** | Progranulin |
| **B3, B4** | CAPG | **D17, D18** | HO-1 | **G7, G8** | Prolactin |
| **B5, B6** | Carbonic Anhydrase IX | **D19, D20** | ICAM-1 | **G9, G10** | Prostasin |
| **B7, B8** | Cathepsin B | **D21, D22** | IL2ra | **G11, G12** | E-Selectin |
| **B9, B10** | Cathepsin D | **D23, D24** | IL6 | **G13, G14** | SERPINB5 (Maspin) |
| **B11, B12** | Cathepsin S | **E1, E2** | IL8 | **G15, G16** | SERPINE1 (PAI-1) |
| **B13, B14** | CEACAM-5 | **E3, E4** | IL18 BPa | **G17, G18** | SNAIL |
| **B15, B16** | Decorin | **E5, E6** | Kallikrein 3 | **G19, G20** | SPARC |
| **B17, B18** | DKK1 | **E7, E8** | Kallikrein 5 | **G21, G22** | Survivin |

| | | | | | |
|---|---|---|---|---|---|
| **B19, B20** | DLL1 | **E9, E10** | Kallikrein 6 | **G23, G24** | Tenascin C |
| **B21, B22** | HER1 | **E11, E12** | Leptin | **H1, H2** | Thrombospondin 1 |
| **B23, B24** | - | **E13, E14** | Lumican | **H3, H4** | TIE2 |
| **C1, C2** | - | **E15, E16** | CCL2/MCP1 | **H5, H6** | u-Plasminogen activator (uPA |
| **C3, C4** | Endoglin | **E17, E18** | CCL8/MCP2 | **H7, H8** | VCAM-1 |
| **C5, C6** | Endostatin | **E19, E20** | CCL7/MCP3 | **H9, H10** | VEGF |
| **C7, C8** | Enolase 2 | **E21, E22** | M-CSF | **H11, H12** | Vimentin |
| **C9, C10** | eNOS/NOS3 | **E23, E24** | Mesothelin | **I1, I2** | Reference Spots |
| **C11, C12** | EpCAM | **F1, F2** | CCL3/MIP1α | **I23, I24** | Negative Control |
| **C13, C14** | ERα | **F3, F4** | CCL20/MIP3α | | |

**Supplementary Movie 1:** Movie of MDA-MB-231 cells seeded onto model group HH ECM. Cells were seeded onto the decellularized ECM and incubated for one day before imaging every 15 minutes over the course of 10 hours.

**Supplementary Movie 2:** Movie of MDA-MB-231 cells seeded onto model group HL ECM. Cells were seeded onto the decellularized ECM and incubated for one day before imaging every 15 minutes over the course of 10 hours.

**Supplementary Movie 3:** Movie of MDA-MB-231 cells seeded onto model group LH ECM. Cells were seeded onto the decellularized ECM and incubated for one day before imaging every 15 minutes over the course of 10 hours.

**Supplementary Movie 4:** Movie of MDA-MB-231 cells seeded onto model group LL ECM. Cells were seeded onto the decellularized ECM and incubated for one day before imaging every 15 minutes over the course of 10 hours.

**Supplementary Movie 5:** Movie of KTB-21 cells seeded onto model group HH ECM. Cells were seeded onto the decellularized ECM and incubated for one day before imaging every 15 minutes over the course of 10 hours

**Supplementary Movie 6:** Movie of KTB-21 cells seeded onto model group HL ECM. Cells were seeded onto the decellularized ECM and incubated for one day before imaging every 15 minutes over the course of 10 hours

**Supplementary Movie 7:** Movie of KTB-21 cells seeded onto model group LH ECM. Cells were seeded onto the decellularized ECM and incubated for one day before imaging every 15 minutes over the course of 10 hours

**Supplementary Movie 8:** Movie of KTB-21 cells seeded onto model group LL ECM. Cells were seeded onto the decellularized ECM and incubated for one day before imaging every 15 minutes over the course of 10 hours.

**Supplementary Movie 9:** Movie of MDA-MB-231 cells seeded onto stretched model group HH ECM. Cells were seeded onto the decellularized ECM and incubated for one day before imaging every 15 minutes over the course of 10 hours

**Supplementary Movie 10:** Movie of MDA-MB-231 cells seeded onto stretched model group HL ECM. Cells were seeded onto the decellularized ECM and incubated for one day before imaging every 15 minutes over the course of 10 hours

**Supplementary Movie 11:** Movie of MDA-MB-231 cells seeded onto stretched model group LH ECM. Cells were seeded onto the decellularized ECM and incubated for one day before imaging every 15 minutes over the course of 10 hours

**Supplementary Movie 12:** Movie of MDA-MB-231 cells seeded onto stretched model group LL ECM. Cells were seeded onto the decellularized ECM and incubated for one day before imaging every 15 minutes over the course of 10 hours

**Supplementary Movie 13:** Movie of KTB-21 cells seeded onto stretched model group HH ECM. Cells were seeded onto the decellularized ECM and incubated for one day before imaging every 15 minutes over the course of 10 hours

**Supplementary Movie 14:** Movie of KTB-21 cells seeded onto stretched model group HL ECM. Cells were seeded onto the decellularized ECM and incubated for one day before imaging every 15 minutes over the course of 10 hours

**Supplementary Movie 15:** Movie of KTB-21 cells seeded onto stretched model group LH ECM. Cells were seeded onto the decellularized ECM and incubated for one day before imaging every 15 minutes over the course of 10 hours

**Supplementary Movie 16:** Movie of KTB-21 cells seeded onto stretched model group LL ECM. Cells were seeded onto the decellularized ECM and incubated for one day before imaging every 15 minutes over the course of 10 hours

**Supplementary Movie 17:** Movie of MDA-MB-231 cells seeded onto mouse mammary fibroblast produced ECM generated with 0 ng/mL leptin within the fibroblast media. Cells were seeded onto the decellularized ECM and incubated for one day before imaging every 15 minutes over the course of 10 hours

**Supplementary Movie 18:** Movie of MDA-MB-231 cells seeded onto mouse mammary fibroblast produced ECM generated with 20 ng/mL leptin within the fibroblast media. Cells were seeded onto the decellularized ECM and incubated for one day before imaging every 15 minutes over the course of 10 hours

**Supplementary Movie 19:** Movie of MDA-MB-231 cells seeded onto mouse mammary fibroblast produced ECM generated with 30 ng/mL leptin within the fibroblast media. Cells were seeded onto the decellularized ECM and incubated for one day before imaging every 15 minutes over the course of 10 hours

**Supplementary Movie 20:** Movie of MDA-MB-231 cells seeded onto mouse mammary fibroblast produced ECM generated with 40 ng/mL leptin within the fibroblast media. Cells were seeded onto the decellularized ECM and incubated for one day before imaging every 15 minutes over the course of 10 hours
